# A comprehensive transcriptome signature of murine hematopoietic stem cell aging

**DOI:** 10.1101/2020.08.10.244434

**Authors:** Arthur Flohr Svendsen, Daozheng Yang, KyungMok Kim, Seka Lazare, Natalia Skinder, Erik Zwart, Anna Mura-Meszaros, Albertina Ausema, Björn von Eyss, Gerald de Haan, Leonid Bystrykh

## Abstract

We surveyed 16 published and unpublished data sets to determine whether a consistent pattern of transcriptional deregulation in aging murine hematopoietic stem cells (HSC) exists. Despite substantial heterogeneity between individual studies, we uncovered a core and robust HSC aging signature. We detected increased transcriptional activation in aged HSCs, further confirmed by chromatin accessibility analysis. Unexpectedly, using two independent computational approaches, we established that deregulated aging genes consist largely of membrane-associated transcripts, including many cell surface molecules previously not associated with HSC biology. We show that *Selp*, the most consistent deregulated gene, is not merely a marker for aged HSCs but is associated with HSC functional decline. Additionally, single-cell transcriptomics analysis revealed increased heterogeneity of the aged HSC pool. We identify the presence of transcriptionally “young-like” HSCs in aged bone marrow. We share our results as an online resource and demonstrate its utility by confirming that exposure to sympathomimetics, and deletion of Dnmt3a/b, molecularly resembles HSC rejuvenation or aging, respectively.

**Key Points:**

1. A comprehensive transcriptome analysis of aged murine hematopoietic stem cells identifies an aging signature;
2. The HSC aging signature is highly enriched for cell membrane-related transcripts and identifies age-associated heterogeneity.

## Introduction

Aging is a natural and time-dependent process, leading to the functional decline of many cell types and tissues, including those of the hematopoietic system. Aged hematopoietic stem cells (HSCs) have less regenerative potential and their overall contribution to formation of mature blood populations, at the clonal level, is restricted and lineage-skewed when compared to their young counterparts – suggesting stem cell exhaustion, one of the hallmarks of aging^1–9^. Transplantation of aged HSCs into young recipients does not improve HSC function^3,5^, indicating that cell-intrinsic aspects play a pivotal role in HSC aging. The differential expression of genes is a prominent read-out of multiple cell-intrinsic factors, and various studies have compared the transcriptomes of young and aged HSCs^10,11,20–24,12–19^. These studies have suggested a plethora of distinct aging-associated mechanisms, including aberrant regulation of genes involved in lymphoid and platelet differentiation^14,22^; elevated inflammation and stress responses^12,17^; chromatin remodeling/DNA methylation^12,23–25^; changes in cell cycle and maintenance of quiescence^15,16,19^; and age-associated replicative stress in HSCs^13^. It remains unclear whether all suggested factors equally contribute to HSC aging, and why different studies report different mechanisms.

Although a substantial number of HSC aging transcriptome studies has been published, no cross-validation of all transcriptomic HSC aging data has yet been carried out. In fact, it is unknown whether a common HSC aging transcriptome exists, and if so, which genes are involved. In order to address this question, we selected 15 studies in which transcriptomes of young and aged murine HSCs were compared, and added to these our own recently published transcriptome analysis^26^. Despite uncovering considerable heterogeneity among different studies, we were able to identify a robust and biologically relevant HSC transcriptomic aging signature. This signature is mainly composed of genes that encode for membrane-associated proteins, many with unknown function(s) in HSC biology. A large fraction of the aging signature genes is upregulated in aged HSCs, indicating transcriptional activation in this population. We also show that aging increases transcriptional heterogeneity among single HSCs and provide evidence that transcriptionally ‘young’ HSCs exist in aged mice. Furthermore, we demonstrate that the top age-associated gene – *Selp* (P-selectin) – contributes to functional decline of HSCs. Finally, we provide an open web-based resource to the community and use independent data sets to demonstrate how our approach can be used to validate novel biological paradigms.

## METHODS

### Mice

All experiments were approved by the Central Commission for animal Testing and by Animal Ethical Committee. Young (4-5 months) and aged (20-24 months) C57BL/6, C57BL/6.SJL (10-14 weeks) and C57BL/6J-kitW-41J/kitW-41J (W41) (originally from European division of Harlan) were obtained from the Central Animal Facility (CDP) at the University Medical Center Groningen. TetOP-H2B-GFP mice provided by Prof. Dr. Rob Coppes (UMCG, Groningen, Netherlands). H2B-GFP expression was induced by providing mice (6 weeks) with food containing doxycycline (Dox) for 6 weeks. Mice were then fed with normal food for 14 weeks. Mice were kept in group cages and housed under conventional in a temperature and day cycle controlled conditions.

### Flow Cytometry

Bone marrow was isolated from the tibia, femur, pelvis, sternum and spine by crushing, and red blood cells were lysed with erylysis buffer. All samples were analyzed on BD FACSCanto™ II (BD Biosciences) and sorted on MoFlo Astrios or XDP cell sorters (Beckman Coulter). Detailed information of the antibodies used for isolation and identifications can be found in supplementary text.

### Retroviral constructs and transduction

*Selp* was amplified from C57BL/6J LT-HSC cDNA and then cloned into pSF91– IRES–GFP vector (originally as pSF91 from C. Baum, Hannover Medical School, Germany, further modified in our lab). Virus was produced by transfecting Platinum-E retroviral packaging cells (Cell Biolabs) according to the manufacturer’s protocol. Viral supernatant was harvested 48 hours after transfection. LT-HSCs were transduced as previously described^6^. Cells were harvested 20-22 hours after transduction for transplantation.

### BM transplantation

For SELP^low^ and SELP^high^ transplantation, 1000 SELP^low^ and SELP^high^ LT-HSCs were isolated from old CD45.2^+^ C57BL/6 mouse and were transplanted together with 2 million CD45.1^+^ W41 bone marrow into lethally irradiated (9 Gy) CD45.1^+^ C57BL/6.SJL recipients, respectively. For *Selp* overexpression transplantation, EV (SF91–IRES–GFP empty) and SELP OE CD45.2^+^ cells harvested after retroviral transduction were transplanted together with 2 million CD45.1^+^ W41 bone marrow into lethally irradiated CD45.1^+^ C57BL/6.SJL recipients, respectively.

### ATAC-sequencing

25,000 LT-HSCs were sorted from young and aged mice in biological replicates. Library preparation was performed according to Buenrostro *et al.,* 2013^27^. In brief, cells were lysed and tagmented for 1h at 37°C. Samples were then PCR-amplified using provided adapter primers for 11 or 12 cycles. Following, samples were size-selected for fragments of 150-1000 bp. Libraries were pooled and sequenced to obtain 30-50 10^6^ reads using a NextSeq 500. Data are deposited under accession number GSE GSE166674

### Immunofluorescence

Images were acquired on a Leica Sp8 confocal microscope and quantified on Fiji Image J.

### RNA content

4000-6000 LT-HSCs were seeded onto spots on an adhesion immunofluorescent slide (VWR). For RNA staining, cells were fixed with 100% methanol for 10 minutes and washed with PBS. Cells were subsequently stained with 1:10000 SYTO RNA Select Green Fluorescent Stain (ThermoFisher) for 20 minutes at room temperature. After washing, coverslips were mounted with ProLong Diamond Antifade Mountant with DAPI.

### RNA pol II

For RNA-Polymerase II staining, cells were fixed and permeabilized with Fixation/Permeabilization Solution Buffer (BD Bioscience) for 20 minutes on ice. After washing, cells were blocked first with Endogenous Biotin-Blocking Kit (Thermo Fisher) followed by 4% BSA for 30 minutes. Cells were stained with 1:100 biotin mouse monoclonal RNA Polymerase II antibody (Novus Bio) at 4°C overnight. Cells were then washed with 0.1% Triton-X-100 and stained with 1:1000 streptavidin Alexa-488 secondary antibody. After washing, coverslips were mounted with ProLong Diamond Antifade Mountant with DAPI.

### Further information

More detailed information about the methods can be found in the supplementary text.

### Additional Resources

Aging signature WEB tool site: http://agingsignature.webhosting.rug.nl/ GitHub with all resources: https://github.com/LeonidBystrykh/data-for-manuscript (will be accessible after manuscript acceptance)

## Results

### Systematic collection of publications reporting gene expression changes in aged HSC

We took advantage of PubMed’s API E-utility to select all informative studies on HSC aging in C57BL/6 mice **(Figure 1A, Supplemental text)**. This search resulted in 16 published studies that met all criteria, and included our own recently published data set^26^ **(Figure 1B/Table S1)**. The various studies used different sequencing platforms, ranging from microarray hybridization, bulk RNA-sequencing (bRNA-seq) to single-cell RNA-sequencing (scRNA-seq)**(Figure 1B/Table S1)** and included mainly 2 age groups **(Figure 1B/S1A)**. We employed two alternative approaches to perform the analysis. First, we compared the self-reported, pre-analyzed lists of DEGs from the selected manuscripts and referred to this as ‘meta-analysis’. In parallel, we re-analyzed all available raw data in the most uniform way; these are referred to as ‘re-analysis’ **(Figure 1A**/**Table S1)**.

**Figure 1:**
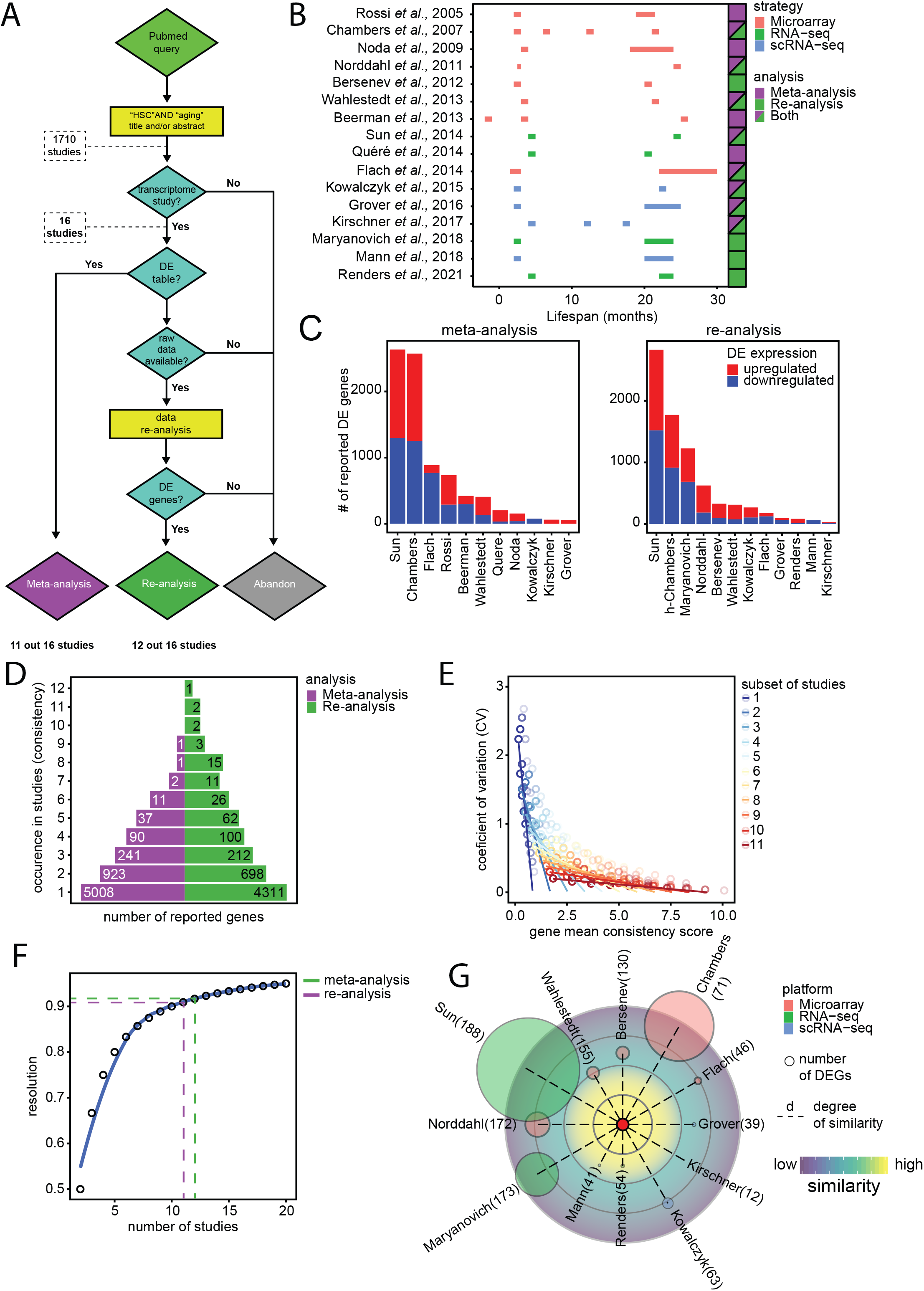
Systematic analysis reveals heterogeneity in transcriptomic studies of young and aged murine HSCs. A) Selection of young and aged HSC transcriptomic studies. Representation of decision tree used to acquire and analyze the different publications used for the systematic analysis. B) Overview of the age groups used in each of the selected studies. The age group of mice used in each study (y-axis) is represented in months (x-axis). Color codes refer to the transcriptome platform used. Squares next to each study indicate in which kind of analysis each study was included. C) Heterogeneity across different transcriptomic studies. Number of differentially expressed genes (DEGs) reported by each study. Left panel shows the number of DEGs as reported by the authors in their respective manuscripts. Right panel shows the number of DEGs identified by us using a uniform re-analysis. Red bars indicate the number of genes that are upregulated upon aging, while blue bars indicate downregulated genes. In the re-analysis panel, “h-Chambers” indicates half-Chambers meaning that just half of the DEGs were taken into account. D) Ranking of DEGs across different studies. Horizontal bars represent the number of DEGs (x-axis, the actual number of DEG genes are shown in each bar), ranked by the number of studies in which they were found (occurrence in studies (consistency), y-axis). Purple and green bars indicate meta-analysis and re-analysis, respectively. E) The list of genes gets more consistent with the increase number of studies added. Simulation of different list assembly using combinations of subsets of 12 studies (gradient color from blue to red). As the number of studies increased, the coefficient of variation (CV, y-axis) decreases. Dots represent the different values and mean consistency for individual genes **(see supplemental text for further details).** F) Predicted resolution to discover consistently reported genes. The model demonstrates that >90% of all HSC aging genes have been identified (y-axis) in the two analytic approaches. Purple and green dashed lines indicate meta-analysis and re-analysis, respectively. G) Extent of overlap between all independent studies and the HSC aging signature (centered red circle, AS). The size of the circles represents the number of DEGs identified in each study; the number of overlapping DEGs with the aging signature is represented next to each study identification; the distance from the center represents the degree of similarity of each study to the HSC aging signature. Studies that show a higher degree of similarity are closer to the AS.

### Individual studies identify highly variable numbers of DEGs

Strikingly, whereas some studies reported thousands of genes to be differentially expressed, others only reported dozens **(Figure 1C)**. When DEGs from the meta-analysis and re-analysis approaches were compared, both demonstrated a significant correlation between fold-change (FC) expression values, indicating that re-analysis of the raw data did not deviate drastically from the originally reported data sets **(Figure S1C).** The observed study-to-study variations might result from differences in mouse gender, group size, exact age, HSC immunophenotyping, intrinsic differences between expression platforms, or unknown environmental effects, including differences in mouse facilities or breeding companies from which the mice originated **(Table S1)**. Principal component analysis (PCA) performed in the re-analyzed sets demonstrated that studies clustered according to their sequencing platform **(Figure S1B)**, although degree of their separation was minor. Overall, this analysis demonstrates that the variability between studies partially has a technical origin, but the discrepancy between studies is not fully explained.

### Most reported age-associated HSC genes have low reproducibility

Collectively these 16 studies reported more than 6000 genes being differentially expressed between young and aged HSCs. Surprisingly, the vast majority of reported genes demonstrated poor reproducibility as almost 80% of all putative age-associated genes were reported in a single study only and not in any other **(Figure S1D)**. Moreover, pair-wise analysis of all different studies showed that some studies had little to no overlapping genes with any other study **(Figure S1E/F).**

### Identification of a HSC aging signature

Despite limited reproducibility between individual data sets, we wondered whether it would be possible to identify consistently DEGs across all studies. To this end, we ranked all reported genes by their occurrence in different studies (consistency) **(Figure 1D)**. This revealed a limited number of highly consistent genes that were frequently documented to be differentially expressed upon HSC aging. The single most consistently deregulated gene was *Selp*. We provide a complete gene list in **Table S2**. We restricted further analysis to genes which were consistently reported in ≥ 4 studies, resulting in a list of 142 genes for the meta-analysis and 220 genes for the re-analyzed data sets. We refer to this as the “aging signature” (AS) **(Figure S1G/ Table S2)**.

### The aging signature is robust and provides higher resolution than individual studies

In order to assess the minimal number of publications required to generate a robust list of consistent aging genes, we modelled all possible combinations of most consistent genes from the re-analyzed data, starting from 1 data set at a time to all combinations of 11 out of 12 data sets **(Figure 2B)**. This analysis generated multiple consistencies for the same genes across different permutations. When the analysis was performed with a small number of data sets, we observed a high degree of variation, which started to diminish when the number of data sets increased. When >6 data sets were used, the number and consistency of genes stabilized **(Figure 1E/ supplemental text)**. Complementarily, we also modeled the predicted resolution – the number of aging genes found – for the same range of studies **(Figure 1F)**. The model predicted that the number of studies that we used for our analyses (11 for meta-analysis and 12 for re-analysis) have high resolution; with the current number of included studies, >90% of all the possible age-associated genes have been found. This also implies that the gene collection in the HSC AS remains stable and addition of novel HSC transcriptomes is unlikely to significantly alter the aging signature.

**Figure 2:**
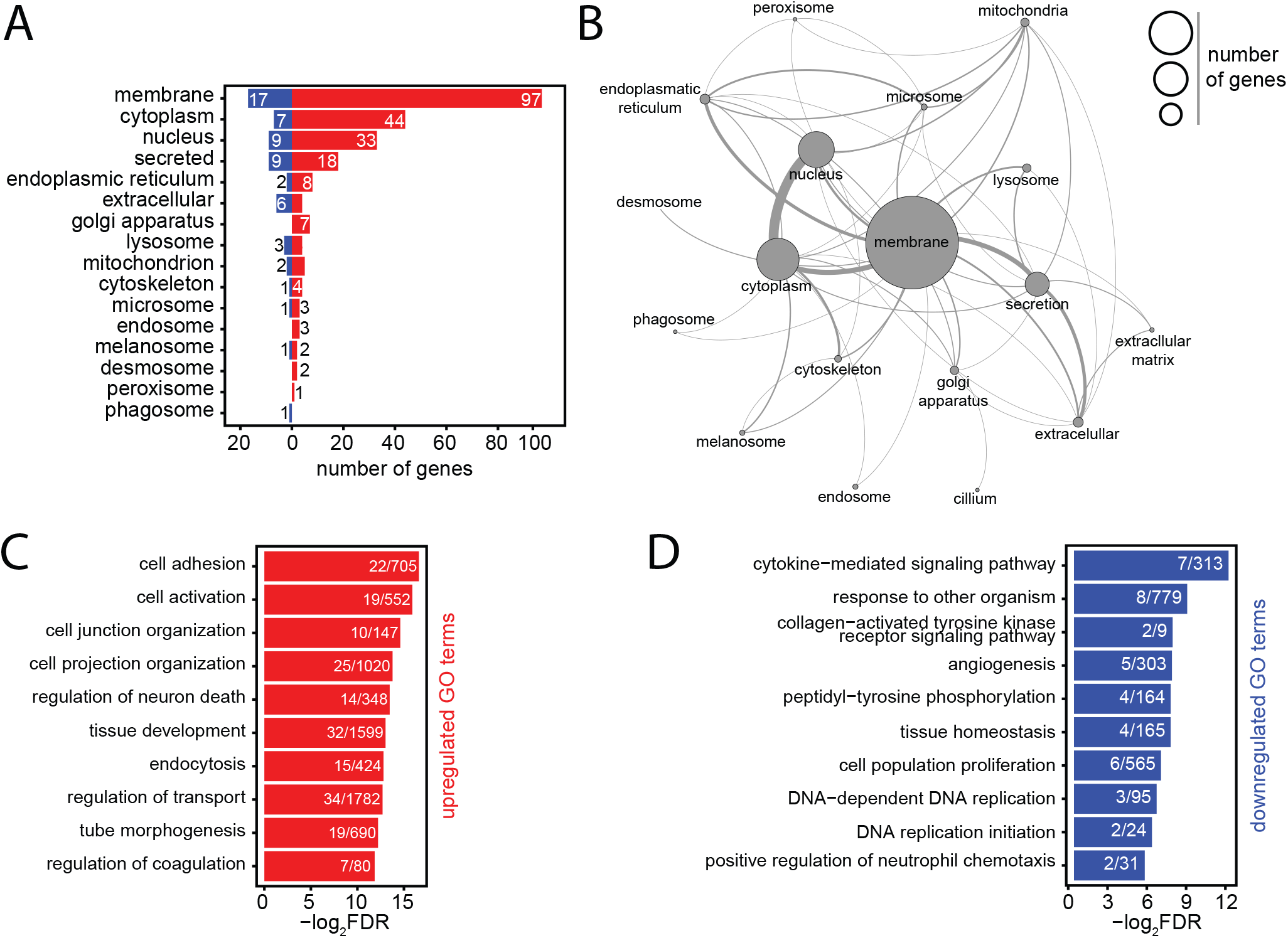
The HSC aging signature is highly enriched for membrane-associated genes. A) Classification of the proteins of the aging signature by cellular compartment. The localization of each protein was determined using UNIPROT classification. Colors of the bars represent upregulation (red) or repression (blue) of genes in the AS. Numbers next to the bars indicate the frequency of genes found in each category. B) Network representation of the aging genes based on the cellular compartment where the encoded proteins are predicted to be localized. The size of each circle represents the number of genes found in each category. Connections between each cellular compartment reflect the number of genes which are shared per cellular compartment. C/D) GO analysis of biological process for AS genes. Horizontal bar represent individual GO terms (y-axis) and their FDR values (x-axis). Numbers in each bar demonstrate the number of genes found per GO term. Red (panel C) and blue (panel D) bars represent GO terms found to be significantly enriched in upregulated and downregulated AS genes, respectively. GO terms are selected based on an FDR < 0.05 cut-off.

We evaluated the degree of similarity between the AS and the individual studies from which it is derived **(Figure 1G)**. All studies demonstrated approximately the same similarity to the AS, with no clear outliers. No single study encompassed all aging genes – but collectively they contribute to the robustness of the AS. We conclude that the AS can be used as a reference for DEGs upon HSC aging.

### The aging signature reveals high enrichment for membrane-related processes in HSC transcriptional activation

We restricted subsequent analyses to the aging signature derived from the re-analyzed data and observed several remarkable features. First, approximately half of the aging genes encode for membrane-associated proteins. **(Figures 2A/B)**. These included many cell adhesion molecules or channels (*Selp*, *Alcam*, *Cd34, Dsg2, Itgb3, Clca3a1*) and cytokine receptors (*Csf2rb, Ebi3, Flt3, Ghr, Il1rapl2, Osmr),* many of which have not been previously associated with HSC biology.

Gene Ontology (GO) analysis identified a total of 46 enriched GO function terms **(Table S3)**. As expected, the most significant GO terms revolve around membrane associated functions **(Figure S2A)**. Among upregulated processes, we observed significant enrichment of different GO terms associated with cell adhesion and activation, and the organization of cell junction and projection (**Figure 2C)**. Within the downregulated processes, we identified significant enrichment for DNA replication, cytokine signaling, tissue homeostasis, and cell proliferation **(Figure 2D)**. Pathway analysis also demonstrated analogous pathways **(Figure S2B)**.

We next collected all associated GO terms associated with every gene present in the aging signature **(Figure S2C)**, resulting in more than 2000 GO terms. The majority of these terms were very specific and contained less than 20 AS genes **(Figure S2C, lower panel)**. We then searched in the AS GO term database for key terms that represent previously described HSC aging mechanisms, including “inflammation”^12^, “replicative stress”^13^, “cell cycle”^15,16^ and “epigenetic drift”^12,23,28^ – and indeed found these. However, the large majority of genes (149) were associated with “membrane” **(Figure S2D)**, denoting that significant changes occur in the HSC membrane upon aging.

Secondly, we observed that a disproportional number (181 genes, ~80%) from the AS were upregulated in aged HSCs **(Figure 3A).** We assessed whether this would be reflected in global transcriptional activation. Comparing young and aged LT-HSCs (LSK CD48^−^ CD150^+^) **(Figure S3A)**, we observed a ~3-fold higher overall RNA yield per cell in aged HSCs **(Figure S3B).** We confirmed this by intracellular staining of young and aged LT-HSCs with the RNA-specific dye SYTO **(Figure S3D)**. In order to investigate if increased RNA abundance in aged LT-HSCs was directly linked to increased transcription, we performed intracellular staining for RNA Polymerase II (POL-II) in young and aged HSCs and indeed observed increased RNA POL-II activity in aged cells **(Figure 3B)**.

**Figure 3:**
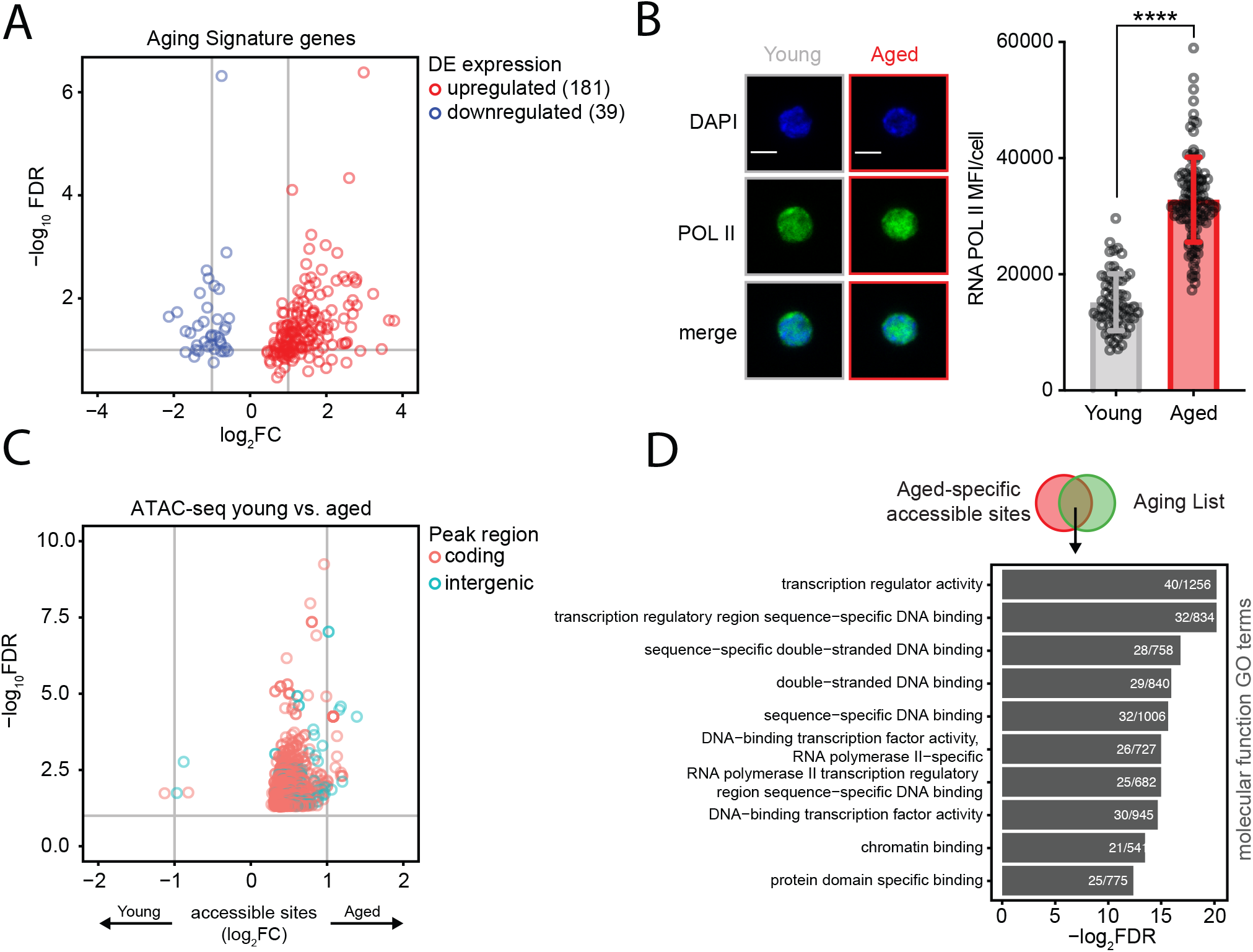
Skewing of HSC aging signature genes reflects transcriptional activation in aged HSCs. A) Skewing of HSC aging signature genes. Volcano plot depicting expression levels of AS genes. 181 upregulated and 39 downregulated genes are represented. Dots represent the average of either adjusted p-values (adjpval, y-axis) and fold-changes (FC, x-axis) per gene in different studies. B) Protein levels of RNA Pol-II in individual young (6 months) and aged (24 months) LT-HSCs. Right panel shows the MFI quantification of young (n = 120 cells) and aged HSCs (n = 139 cells). Scale bars in IF images are 5 μm. ±SD is shown. **** p < 0.0001. C) Aged HSCs demonstrate more age-specific chromatin accessible sites than young HSCs. Volcano plot depicting differentially accessible sites (peaks) between young and old HSCs measured by ATAC-seq. Dot colors demonstrate peaks either in genomic regions overlapping with coding (red) or non-coding (blue) annotations. Peaks with negative FC values are significantly more accessible in young HSCs whereas peaks with positive FCs values are more significantly accessible in aged HSCs. Gene symbols annotate peaks which overlap with gene bodies from some AS genes. D) Transcriptional program is associated with ATAC-seq of young and aged HSCs. GO analysis of functional process for overlapping aged-specific ATAC-seq accessible sites with genes from AL. Horizontal bar represent individual GO terms (y-axis) and their FDR values (x-axis). Numbers in each bar demonstrate the number of genes found per GO term. GO terms are selected based on a FDR < 0.05 cut-off.

Finally, we evaluated the number of chromatin accessible sites by ATAC-seq. In agreement with the data above, aged HSCs showed an increased number of overall accessible sites compared to young HSCs **(Figure 3SD).** In addition, differential analysis of accessible sites demonstrated strong skewing towards more open sites in aged HSCs **(Figure 3C)**. We then speculated that aged-specific accessible sites could contribute to deregulated expression. To investigate this, we overlapped the age-restricted accessible sites with the genes in the aging list – which would reflect genes more accessible in aged HSCs and therefore upregulated – and performed a GO analysis **(Figure 3D)**. This revealed high enrichment for transcription-related processes, further reinforcing our hypothesis that aged HSCs are transcriptionally activated.

Taken together, the aging signature suggests that aged HSCs undergo adaptation at cellular and functional levels through membrane-related processes and that aged HSCs demonstrate higher transcriptional activation.

### The aging signature as a reference for age-associated studies

We aimed to test whether the aging signature can be used as a reference to (re-) analyze existing and future HSC gene expression profiles. To this end, we developed two independent approaches where custom transcriptome sets can be compared to the aging signature.

First, we designed two scoring methods, the discrete Gene Set Enrichment (dGSE) algorithm and a Rank Score (RS), both of which are able to cope with semi-ordered (discrete) lists of genes **(Supplemental text)** and can be used for testing bulk RNA-seq sets. To benchmark both methods, we individually tested the 12 HSC aging transcriptome studies that comprise the aging signature **(Figure 4A)**. As expected, we observed that the different data sets had a wide range of enrichment scores when compared to the aging signature.

**Figure 4:**
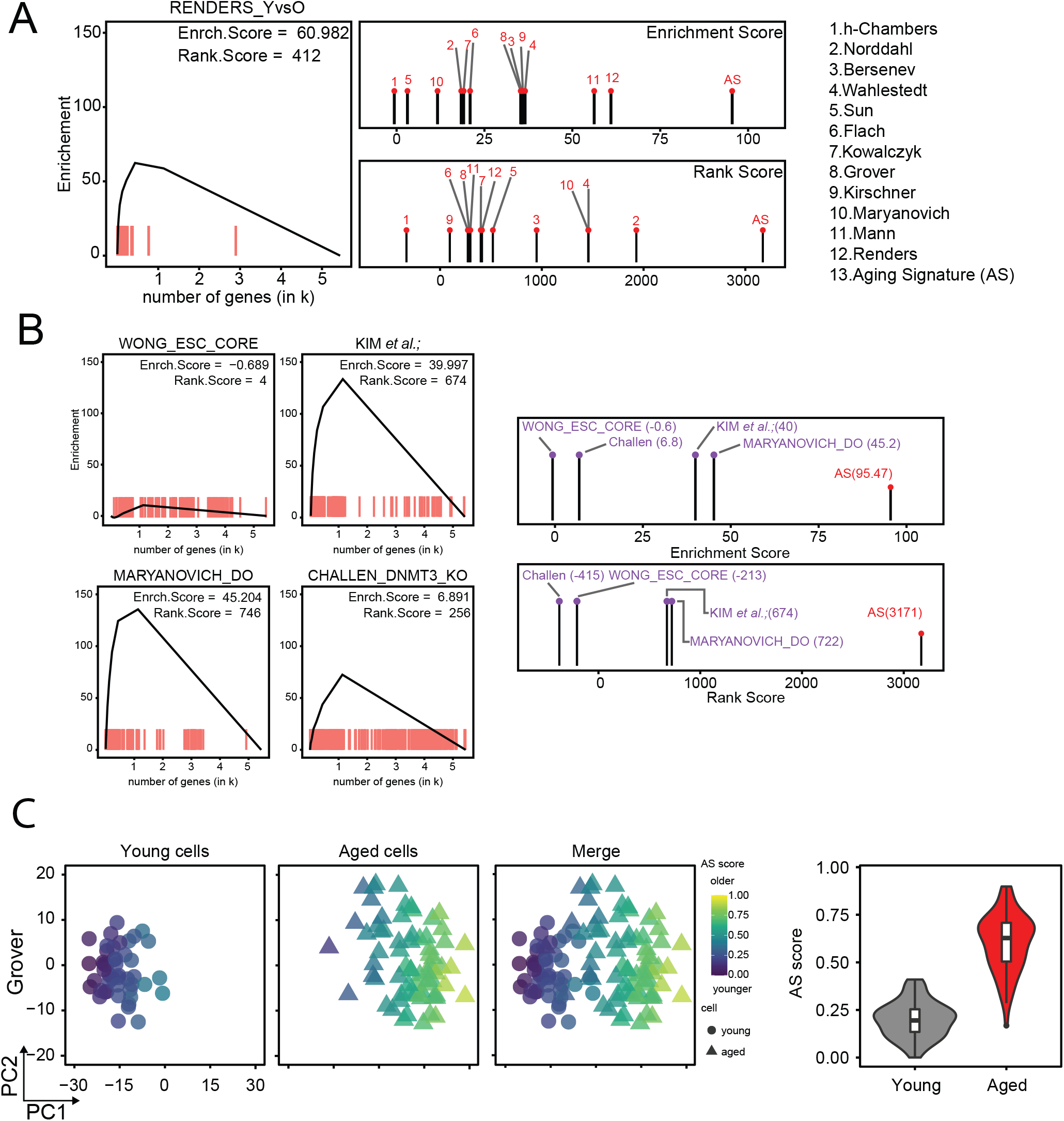
Employing the HSC aging signature as a bona fide transcriptomic reference. A) Using Rank Score (RS) and discrete gene set enrichment (dGSE) analyses to compare existing data sets to the HSC aging signature. Left panel: graphical representation of the dGSE enrichment analyses from our own generated HSC transcriptome data was compared to the HSC aging signature. Right panel: Plots of RS (upper) and dGSE (lower) enrichment analyses of all individual studies that comprise the AS. Numbers refer to the studies shown on the right. B) Searching for an aging signature using RS and dGSE analyses in independent data sets. Upper left panel: Lack of enrichment for aging genes in an embryonic stem cell signature from *Wong et al.* Upper Right panel: Enrichment for the top250 genes (sorted by FDR value) of Kim *et al.;* Lower right panel: Significant enrichment for an HSC aging signature among genes downregulated in “rejuvenated” HSCs from Maryanovich *et al*. Lower Right panel: Significant enrichment for a HSC signature among differentially expressed genes in HSCs isolated from young DNMT3A/B KO mice from Challen *et al.* The right-most panels show combined RS and dGSE scores for each test set. C) Single-cell AS scores. Separation of young (circles) and aged (triangles) single-cell transcriptomics and their respective AS score. From Grover scRNA-seq data (48 young and 68 aged cells), individual cells were separated by PC1 (x-axis) and PC2 (y-axis) and color-coded according to their individual aging signature score. Left column display only young cells; middle column shows just aged cells and right column shows both populations. Cells are colored score from low (blue) to high (yellow) AS scores. On the far-right panel, violin and boxplot demonstrate the overall AS score (y-axis) distribution of the young (grey) and aged (red) cells.

We next used transcriptomic data sets from three unrelated studies. The first independent data set compared young and physiologically aged HSCs (Kim *et al*., manuscript in revision). The second involved HSCs that were “rejuvenated” by a sympathomimetic drug acting on adrenoreceptor β3^18^. The third study used expression data from HSCs that were derived from *Dmnt3a/b* deficient mice that show accelerated aging phenotypes^29^. As a negative control, we included a transcriptomic signature of embryonic stem cells (ES)^30^. As expected, the ES signature demonstrated low scores for aging genes. In contrast, the independent set of young and physiologically aged HSCs demonstrated enrichment when compared to the AS. When we compared the list of genes that were downregulated in “rejuvenated” old HSCs to the AS, we also found strong enrichment. The transcriptome of young HSCs isolated from Dmnt3a/b KO mice also scored high for HSC aging genes. These data suggest that both β3 sympathomimetic signaling and loss of *Dnmt3a/b* molecularly phenocopy normal HSC aging, additionally confirming the value of our method **(Figure 4B)**.

This approach allowed us to investigate the relationship of our HSC aging signature to a previously established multi-organ inter-species transcriptomic signature^31^. Expectedly, we observed a mild enrichment between both signatures, denoting that some more general aging-associated genes are also present in HSC aging **(Figure S4A)**. Genes present in both signatures are related to inflammatory response. We also investigated the involvement of TGF-β signaling, which has been implicated in HSC aging^21,23,32^, in the aging signature. We compared different molecular signatures of TGF-β signaling^33,34^ to our aging signature, but none of the sets demonstrated high scores. Furthermore, only 5 TGF-β target genes from all sets (*Bmpr1a*, *Id2*, *Jun*, *Itgb3* and *Mef2c*) are present in the AS. Thus, TGF-β signaling, from a transcriptional perspective, is not robustly involved in HSC aging. **(Figure S4A)**.

Interestingly, single-cell transplantations have revealed that the overall engraftment potential of aged HSCs is substantially lower than that of young HSCs^2^. However, some aged HSCs display engraftment performance similar to young cells, suggesting that the aged HSC pool is functionally heterogenous. Thus, we wondered whether the AS can be used to also identify intrinsic variation between cells from the same population. To identify such “young-like” cells using single-cell transcriptomic studies, we assigned an Aging Signature score (AS score) to individual cells **(Figures 4C/S4B)**. This revealed that separation of young and aged cells is in good agreement with their assigned AS scores where aged cells score higher and young cells lower, also predicting their age. Interestingly, some aged cells had a low AS score and were close to the young cells cluster, suggesting that these cells are, at least transcriptionally, “young-like”.

Taken together the HSC aging signature demonstrated to be a *bode fine* transcriptional reference to physiological aging and thus a valuable resource for the scientific community. It allowed us to demonstrate that the aged HSC pool is heterogeneous and that some aged cells are transcriptionally similar to young HSCs. We provide both (bulk and single-cell) analyses as a resource to the community at https://agingsignature.webhosting.rug.nl/.

### Machine learning algorithms independently identify the best age-associated gene predictors

Employing machine learning strategies in conjunction with scRNA-seq data have allowed for prediction and detection of different hematopoietic populations^35,36^. We envisioned that by applying this approach to the age-related scRNA-seq data sets would allow for the identification of genes which best divide young and old HSCs. It would also serve as an independent and alternative approach to DE analysis. We trained and tested two different machine learning algorithms – ADAboost and Random Forest (RNDforest) – for identification of the best gene(s) predictors. Interestingly, both algorithms determined that *Nupr1*, *Selp*, and *Sult1a1* were the best predictors **(Figure S5SA)**. We tried to restrict the aging-related list to the best 20 age-associated genes. With those 20 genes, both algorithms performed well on identifying young and aged HSCs **(Figures 5A/B/S5B/C)** when compared to the full list of AS genes. Attempts to further narrow down the age-predictors from 20 to 5 genes, however, showed considerable deterioration in prediction capacity compared to larger lists of age-predictors **(Figure S5D)**. Out of 20 gene predictors, the majority are also membrane-related **(Figure 5C)**.

**Figure 5:**
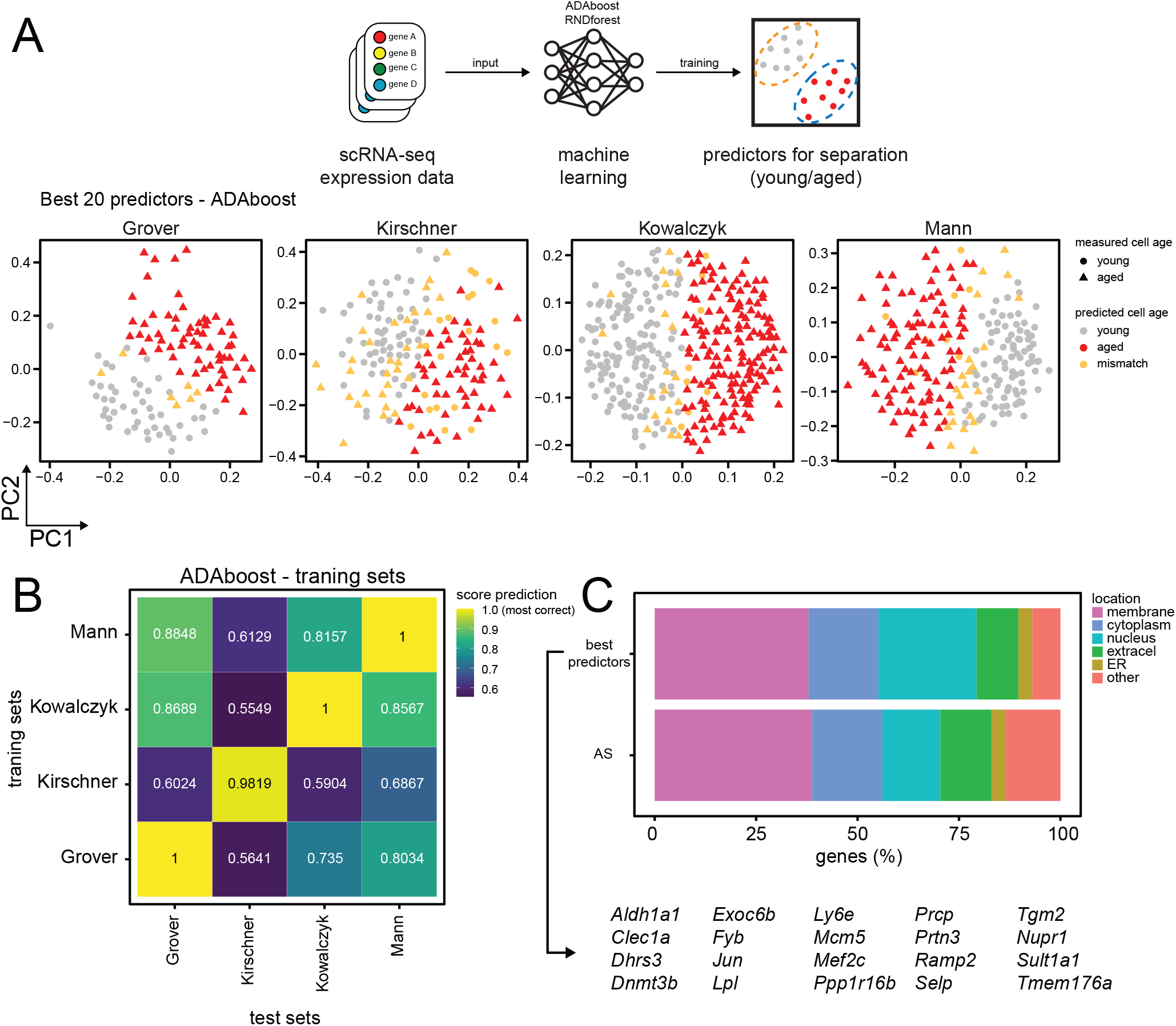
Machine learning identifies best 20 gene predictors for separating young and aged HSCs in scRNA-seq. A) Machine learning is able to predict with high accuracy young and aged HSCs. Top panel depicts a schematic representation of machine learning application into transcriptomic data from different scRNA-seq sets. Lower panel represents the output of one of the algorithm used (ADAboost) in individual cells from different sets. Young (circles) and aged (triangles) single-cell were separated by PC1 (x-axis) and PC2 (y-axis) and color-coded according to the match between the measured aged of the cells and the predicted aged measured by the algorithm (grey for young cells; red for aged cells and orange for mismatched cells). B) Machine learning scores varies depending on which training set is used. Heatmap from ADAboost training depicting different overall scores for different scRNA-seq sets used. The overall score is color-coded from blue (lower scores) to yellow (higher scores). Sets on the x-axis (training sets) were used to train the algorithm and the following sets on the y-axis (test sets) were scored according to training. C) The best machine learning genes predictors have also high enrichment for membrane-associated proteins. Horizontal stacked plot comparing the list of the 20 best predictors extracted from machine learning algorithms (best predictors) and the aging signature genes. The percentage of genes in each cellular location is represented as a percentage (x-axis) and divided by category (different colors). The 20 best predictors gene symbols are represented below.

### The top aging gene – Selp – affects HSC functioning

The most consistent gene from the AS – *Selp* – **(Table 2)** was also among the best machine learning predictors **(Figure 5)**. *Selp*, which encodes for the membrane adhesion molecule P-selectin, was confirmed to be differently expressed also at the protein levels in young and aged hematopoietic progenitor subsets by flow cytometry. **(Figures 6A/S6A)**. To investigate whether Selp affects HSC functioning we competitively transplanted 1000 SELP^low^ or 1000 SELP^high^ aged LT-HSCs **(Figure S6B)** into lethally irradiated young recipients. Whereas overall engraftment levels were similar for SELP^low^ and SELP^high^ LT-HSCs, we found that SELP^high^ LT-HSCs produced more myeloid cells, suggesting that SELP^high^ HSCs are myeloid biased **(Figures 6B/S6C)**.

**Figure 6:**
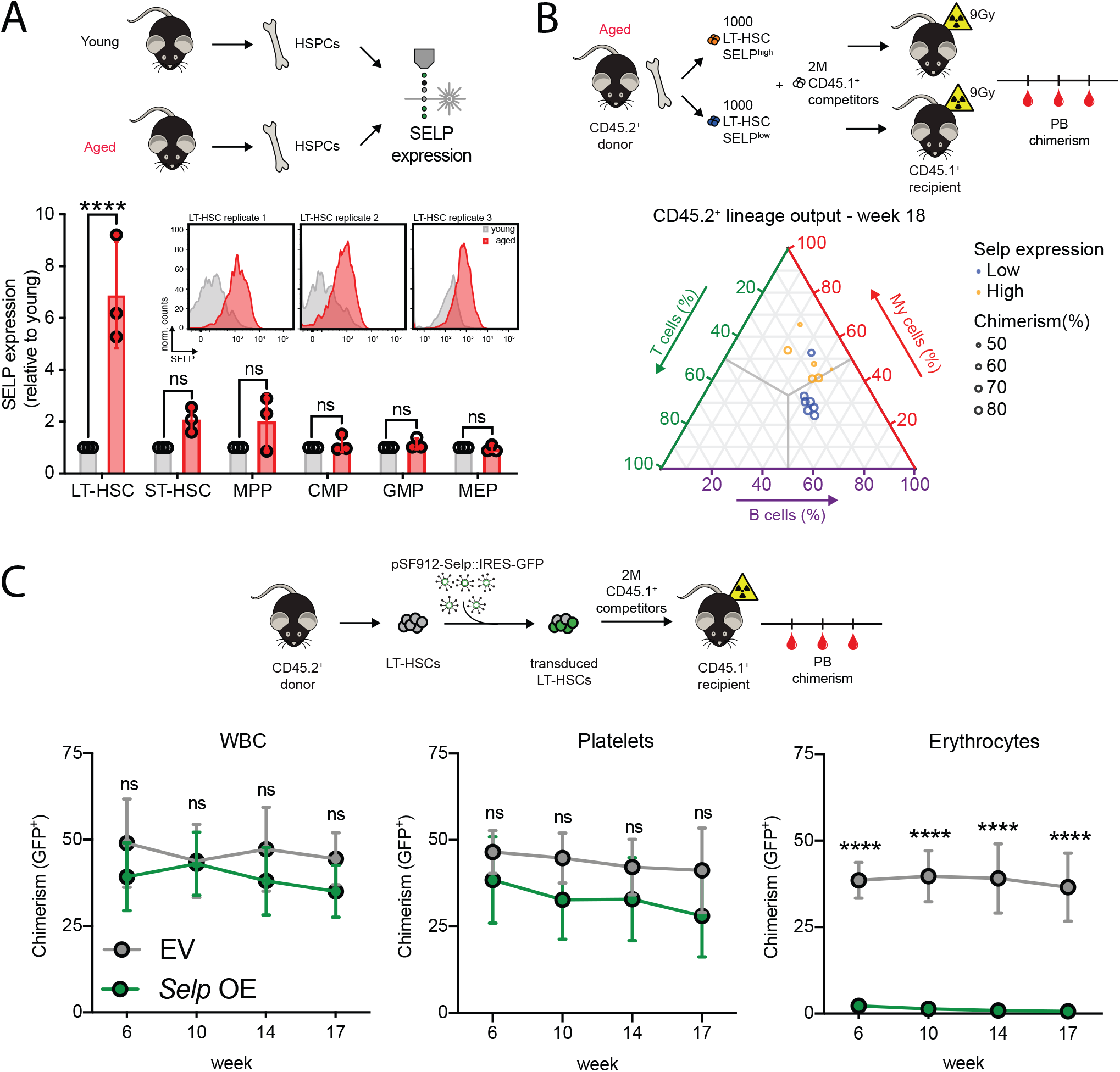
Selp (P-Selectin) is the top aging signature candidate and contributes to HSC functional decline. A) SELP protein expression level across different hematopoietic stem and progenitors (HSPCs) in young (grey) and aged (red) mice. Panel shows fold-changes of aged (n = 3 mice) samples normalized to their young (n = 3 mice) counterparts. Dots show biological replicates while the bars represent the mean values for the given population. Flow cytometry plot for SELP expression in young and aged LT-HSCs population in the 3 biological replicates. B) Selp^high^ LT-HSCs display a myeloid bias. The upper panel shows the experimental approach for competitive transplantation. SELP^low^ (blue) and SELP^high^ (orange) LT-HSCs were purified from aged mice and competitively transplanted (n = 6-7 mice for each group). Overall engraftment (WBC), and T cells levels were similar comparing both groups whereas Myeloid (My) and T-cells levels indicate that transplanted LT-HSC SELP^high^ have Myeloid-bias. C) SELP OE recipients show impairment of erythroid differentiation. The upper panel shows the experimental approach for competitive transplantation of SELP OE and EV (empty vector) HSCs. Lower panel displays peripheral blood (PB) output of the GFP^+^ (transduced cells) of both groups (n = 3 per group). Within the GFP^+^ population, the number of Ter-119^+^ population is significantly reduced. For all panels, ±SD is shown. ns = non-significant; * p < 0.05; ** p < 0.01; n indicates biological replicates

To determine the role of *Selp* in the functioning of LT-HSCs, we overexpressed *Selp* in young LT-HSCs, and competitively transplanted these in young recipients. Whereas *Selp* overexpression did not affect differentiation to leukocytes and platelets, erythroid production was almost completely abolished **(Figure 6C).** We also employed a Tet-On H2B-GFP reporter mouse to determine the dormancy or activation status of SELP^low^ and SELP^high^ LT-HSCs. GFP fluorescence was significantly higher in SELP^low^ LT-HSCs, indicating that myeloid-biased SELP^high^ LT-HSCs were more actively cycling **(Figure S6D).**

Thus, higher levels of SELP in LT-HSCs functionally contribute to myeloid bias, erythroid blockage and HSC activation. Collectively, this demonstrates that SELP is not only a marker of aged HSCs but has the capacity to contribute to the functional impairment of HSCs.

## Discussion

In this study we demonstrate that individual studies have limited resolution to identify genes altered in HSC aging, but collectively they provide an unprecedented profile of transcriptomic alterations in HSC aging **(Figure 1D)**. From this extensive repertoire of studies, we were able to identify a comprehensive, robust, and stable transcriptomic signature of HSC aging, which we validated by 3 independent data sets.

The HSC aging signature is highly enriched for membrane-associated transcripts **(Figure 2A)**. Several of these have been previously associated with aspects of the HSC aging process and thus document the rigor of our approach **(Table 1)**. However, the role of most other genes in HSC (aging) biology remains unknown. The strong involvement of cell surface molecules does provide insight into the long-standing discussion whether HSC aging results from cell-intrinsic or cell-extrinsic molecular alterations. This enrichment of cell membrane-associated transcripts suggests that physiologically aged HSCs communicate differently with their immediate environment compared to their young counterparts.

**Table 1.** Top 10 Aging signature genes with experimental validated function in HSC aging

| Gene Symbol | Consistency (out of 12) | Average Log<sub>2</sub>FC | Function | Phenotype | Reference |
| --- | --- | --- | --- | --- | --- |
| ***Itgb3*** | 9 | 1.25±0.48 | Cell adhesion;<br>Signal transduction | *Itgb3*<sup>high</sup> identifies myeloid-biased HSCs | 17,39 |
| ***Vwf*** | 8 | 1.37±0.66 | Cell adhesion;<br>Signal transduction | *Vwf*<sup>high</sup> identifies myeloid-biased HSCs | 14,40,41 |
| ***Neo1*** | 8 | 1.84±1.18 | Cell adhesion;<br>Signal transduction | *Neo1*<sup>high</sup> identifies more dormant HSCs | 26,42 |
| ***Alcam*** | 8 | 1.39±0.62 | Cell adhesion | *Alcam*<sup>+</sup> identifies myeloid-biased HSCs and with reduced potential | 43 |
| ***CD86*** | 6 | -1.39±0.36 | Signal transduction | Required for lymphoid differentiation | 44 |
| ***Gadd45g*** | 6 | 1.54±0.99 | Stress response | *Gadd45g*<sup>-/-</sup> show increased self-renewal and deficient Mk differentiation | 45 |
| ***Satb1*** | 6 | -1.21±0.50 | Epigenetic effector | Required for lymphoid differentiation | 46 |
| ***Sdpr*** | 6 | 1.88±1.61 | Transcription regulation | Involved in PRC2 expression regulation | 47 |
| ***Slamf1*** | 5 | 0.92±0.16 | Cell adhesion<br>Signal;<br>transduction | *Slamf1*<sup>high</sup> identifies myeloid-biased HSCs and with reduced potential | 48,49 |
| ***Dnmt3b*** | 4 | -1.09±0.62 | Epigenetic regulator | *Dnmt3a/b*<sup>-/-</sup> show differentiation blockage/ increased self-renewal | 29,50,51 |

Our study also provides novel insight into the transcriptional activation state of HSCs (**Figure 3)**. In line with recent reports that loss of heterochromatin leads to age-associated phenotypes in HSCs^37^, RNA-specific quantification, RNA POL-II activity and chromatin accessibility analyses, all indicate that aged HSCs are, at a population level, more transcriptional active than young HSCs.

The aging signature can be employed as a *bona fide* transcriptional reference for physiological HSC aging for both bulk and single-cell transcriptomic studies **(Figure 4)**. We provide protocols and a user-friendly and interactive website – https://agingsignature.webhosting.rug.nl/ – to allow users to execute custom queries and compare new data sets against the aging signature that we present here.

Our approaches also allowed powerful analyses and uncovered novel perspectives into HSC aging from a single-cell perspective. By applying our AS score to single-cell transcriptomic studies, we determined that HSCs isolated from aged mice contain a fraction of cells which behave, at least transcriptionally, like young HSCs **(Figure 4C)**. Further studies are necessary to understand if such cells indeed are functionally superior and also why such cells do not seem to “age”. We also successfully employed machine leaning to further identify best age-associated gene predictors **(Figure 5)**.

Both computational approaches demonstrated that *Selp* expression is strongly associated with HSC aging. Previous data have shown that *Selp*-deficient HSCs display a competitive advantage over wild-type HSCs^38^. Here we document that high levels of SELP expression induced myeloid-biased, erythroid blockage and cell activation, all hallmarks of HSC aging^8,9^ **(Figure 6)**. Together with *Selp,* which is a well-known inflammatory marker, *Nupr1* is one of the top-listed predictors and belongs to a group of inflammatory and stress-responding transcription factors, not earlier detected in relation to the aging phenotype. Although we were able to reduce the list of age-related genes, it is clear that HSC aging cannot be reduced to a single gene. Instead, our machine learning analysis suggests that multiple age-related genes are involved in the HSC aging process. The idea of a single gene explaining most of the aging phenotype sounds unlikely, and that multiple cell surface markers are required to prospectively separate young and aged HSCs.

Nevertheless, our data and others demonstrate that (at least some of) the aging genes that we identify are not merely markers of aged HSCs, but also are involved in age-dependent functional decline. The identification of the HSC aging signature does not resolve whether altered expression is the cause or consequence of loss of HSC functioning during aging. Future functional studies are required to assess whether perturbation of AS genes can improve functioning of aged stem cells. We believe that this resource is valuable in assisting the aging research community to verify involvement of genes in physiological HSC aging.

## Acknowledgements

We thank T. Bijma, G.Mesander and J. Teunis from UMCG Flowcytometry Unit facilities for their assistance on cell sorting; Klaas Sjollema from UMCG Microscopy and Imaging Center (UMIC) for assistance with confocal microscopy; B Bakker, M Gerritsen, E Verovskaya and all members of the Ageing Biology and Stem Cells laboratory for discussions. This work was supported by the Netherlands Organization for Scientific Research/Mouse Clinic for Cancer and Aging, the Landsteiner Foundation for Blood Transfusion Research (LSBR1703), a China Student Council Fellowship awarded to D.Y., Marriage, a EU FP7 Marie Curie Initial Training Network (Contract number 316964) and ARCH, a European Union’s Horizon 2020 research and innovation programme under the Marie Skłodowska-Curie grant agreement No 813091. B.v.E. was supported by grants from the DFG (EY 120/1-1), Else Kröner-Fresenius Foundation (2016_A58) and the German Cancer Aid (Deutsche Krebshilfe; 70113138). The FLI is a member of the Leibniz Association and is financially supported by the Federal Government of Germany and the State of Thuringia.

## Author Contributions

Conceptualization, A.F.S., D.Y., S.S., G.d.H. and L.B.; Methodology, A.F.S., D.Y., and L.B.; Investigation, A.F.S., D.Y., S.S., A.A. K.K; A.M-M., B.v.E. and L.B.; Formal Analysis, A.F.S., D.Y., S.S. K.K; A.M-M., B.v.E. and L.B.; Data Curation, A.F.S., E.Z., L.B.; Resources, E.Z. and A.A.; Writing – Original Draft, A.F.S.; Writing – Review & Editing, A.F.S., D.Y., G.d.H. and L.B.; Supervision, G.d.H. and L.B.

