## Supplemental Figures for "A comprehensive transcriptome signature of murine hematopoietic stem cell aging"

### Supplemental Figure 1

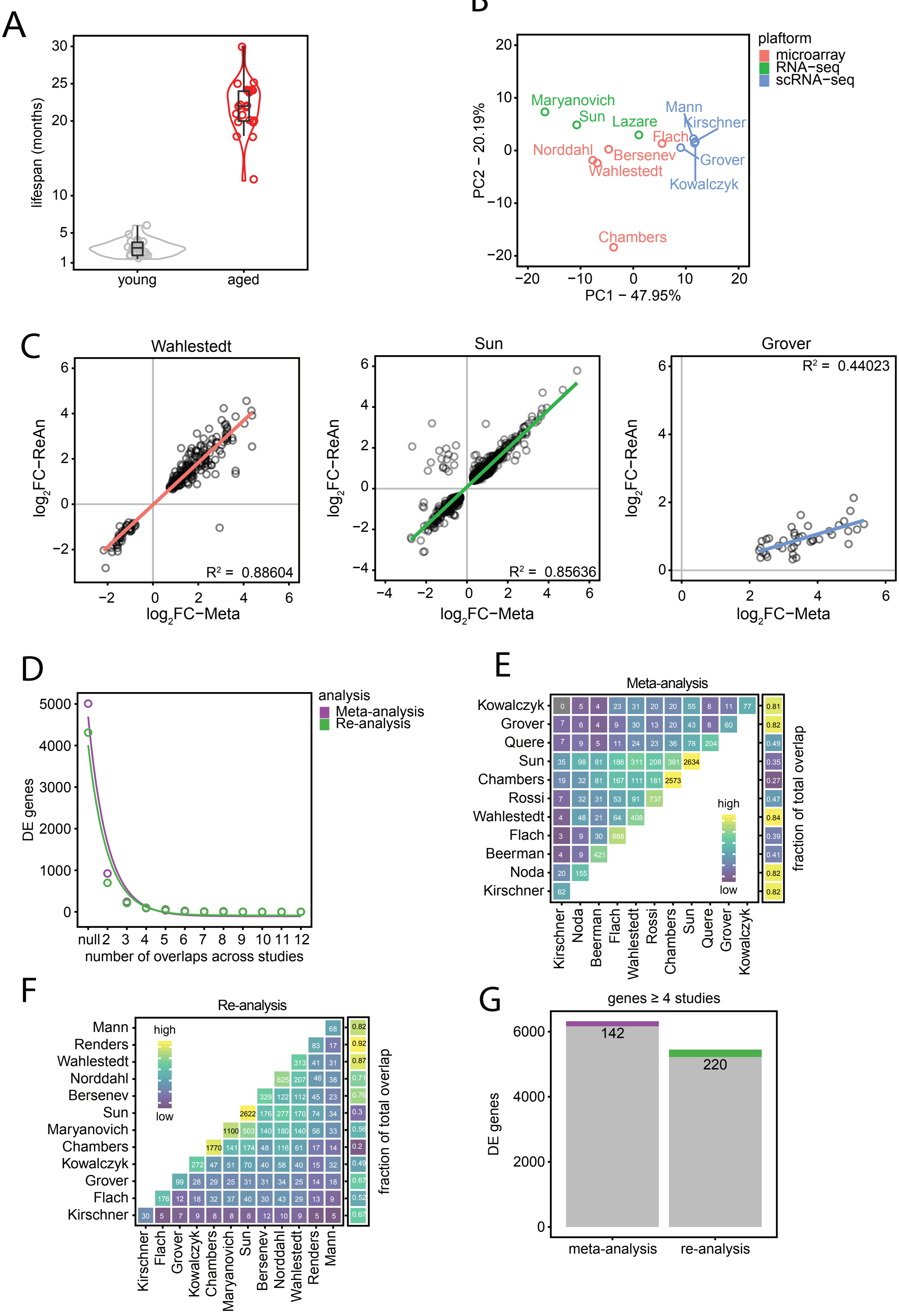

### Supplemental Figure 2

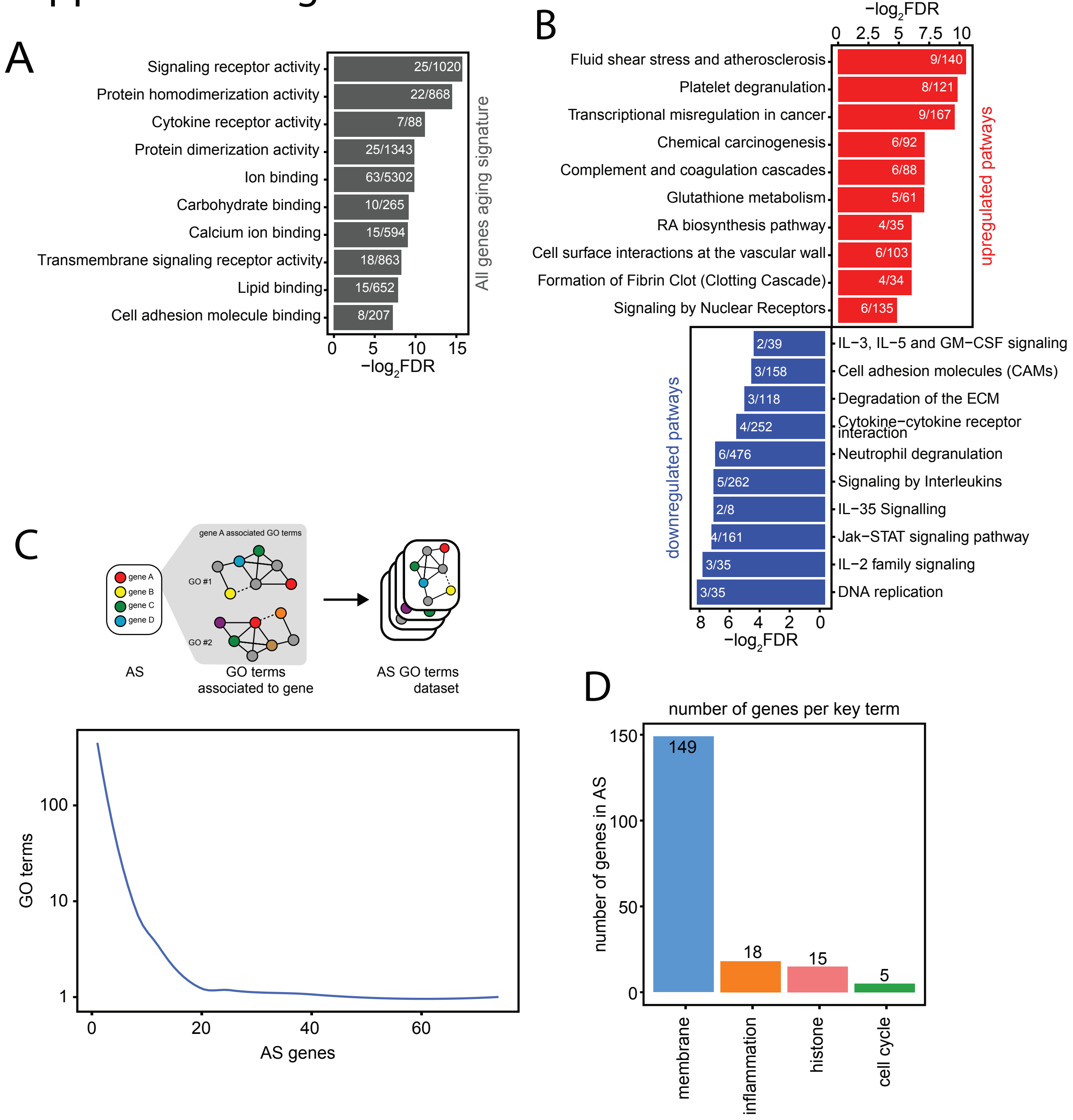

### Supplemental Figure 3

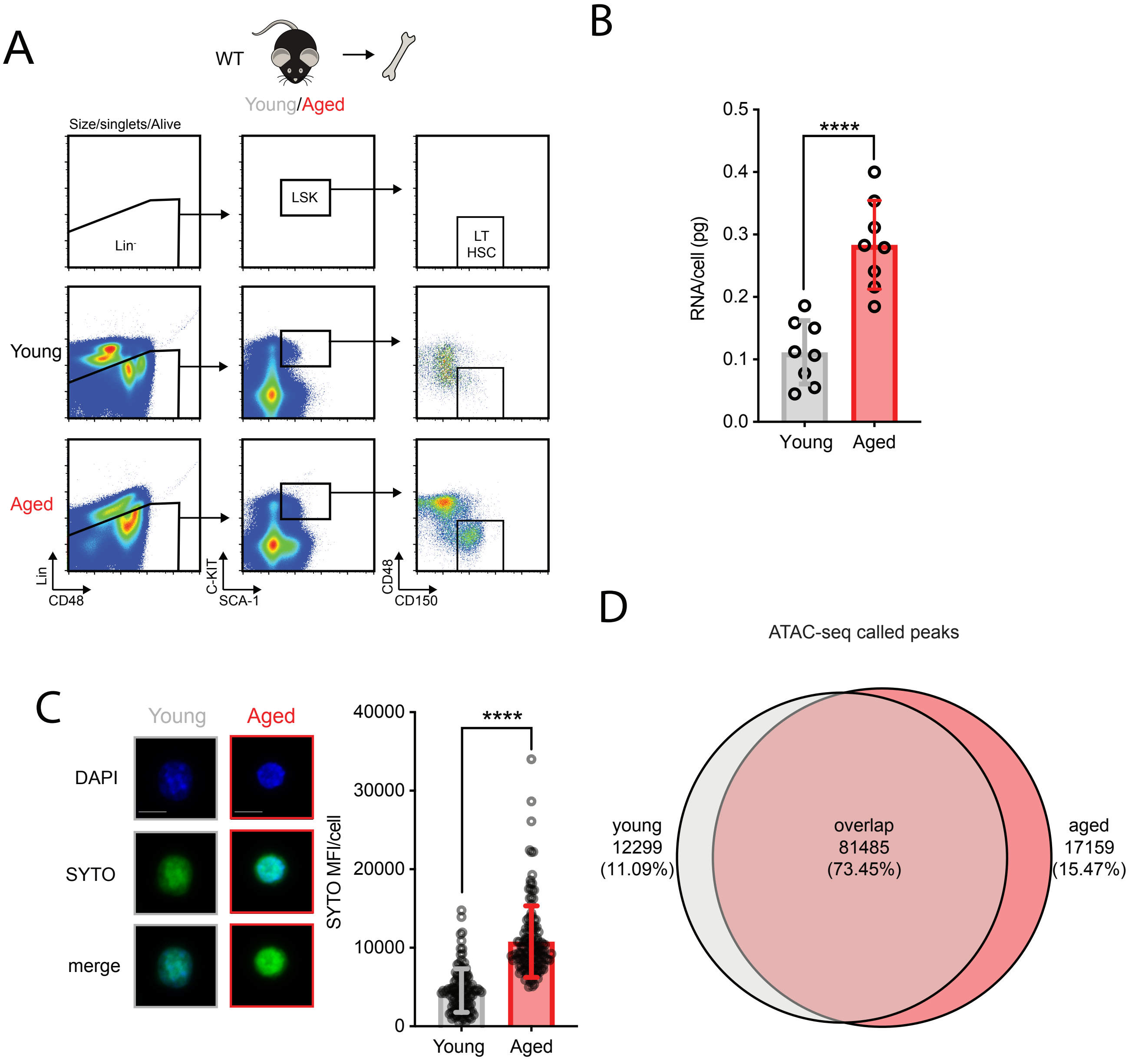

### Supplemental Figure 4

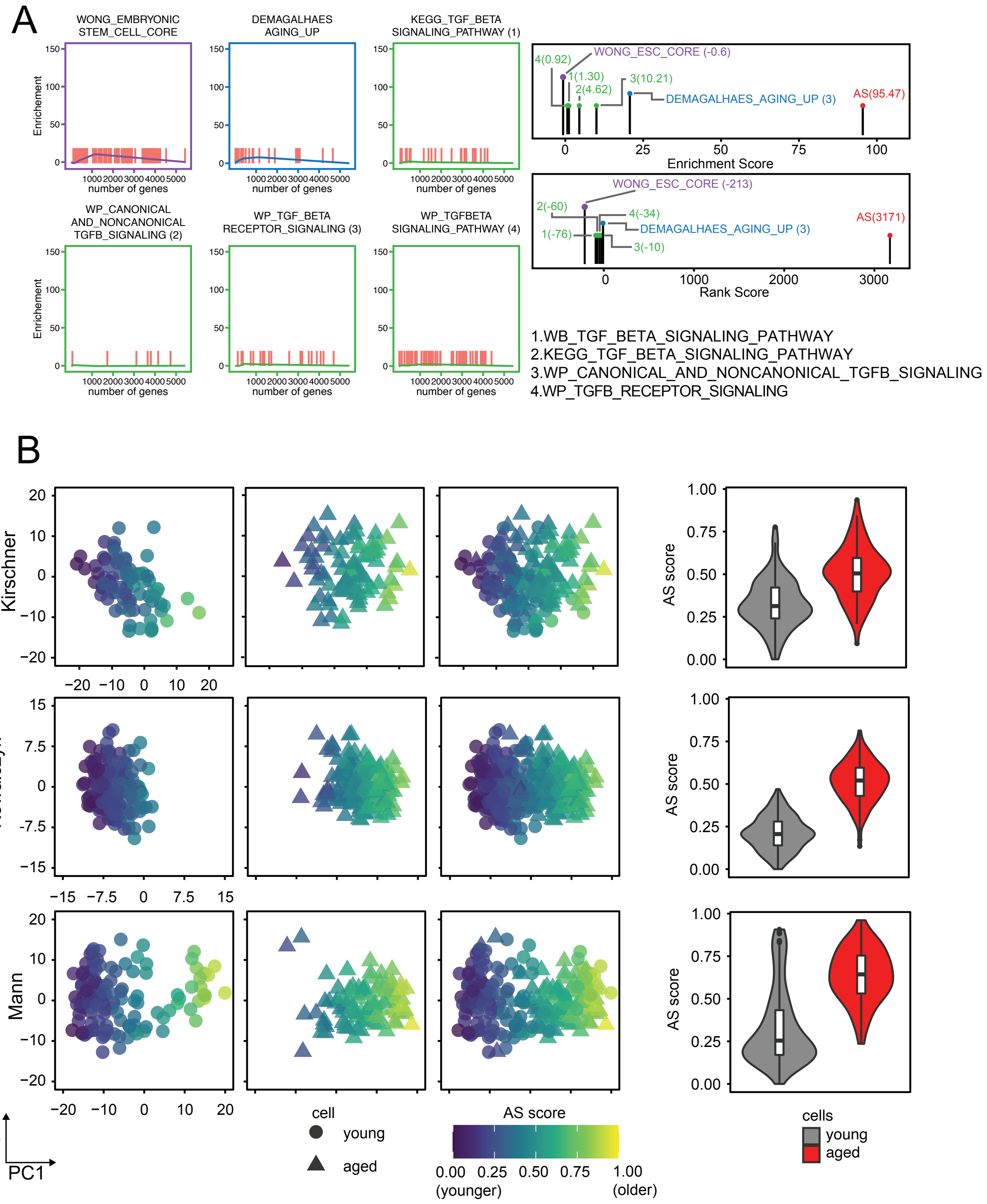

### Supplemental Figure 5

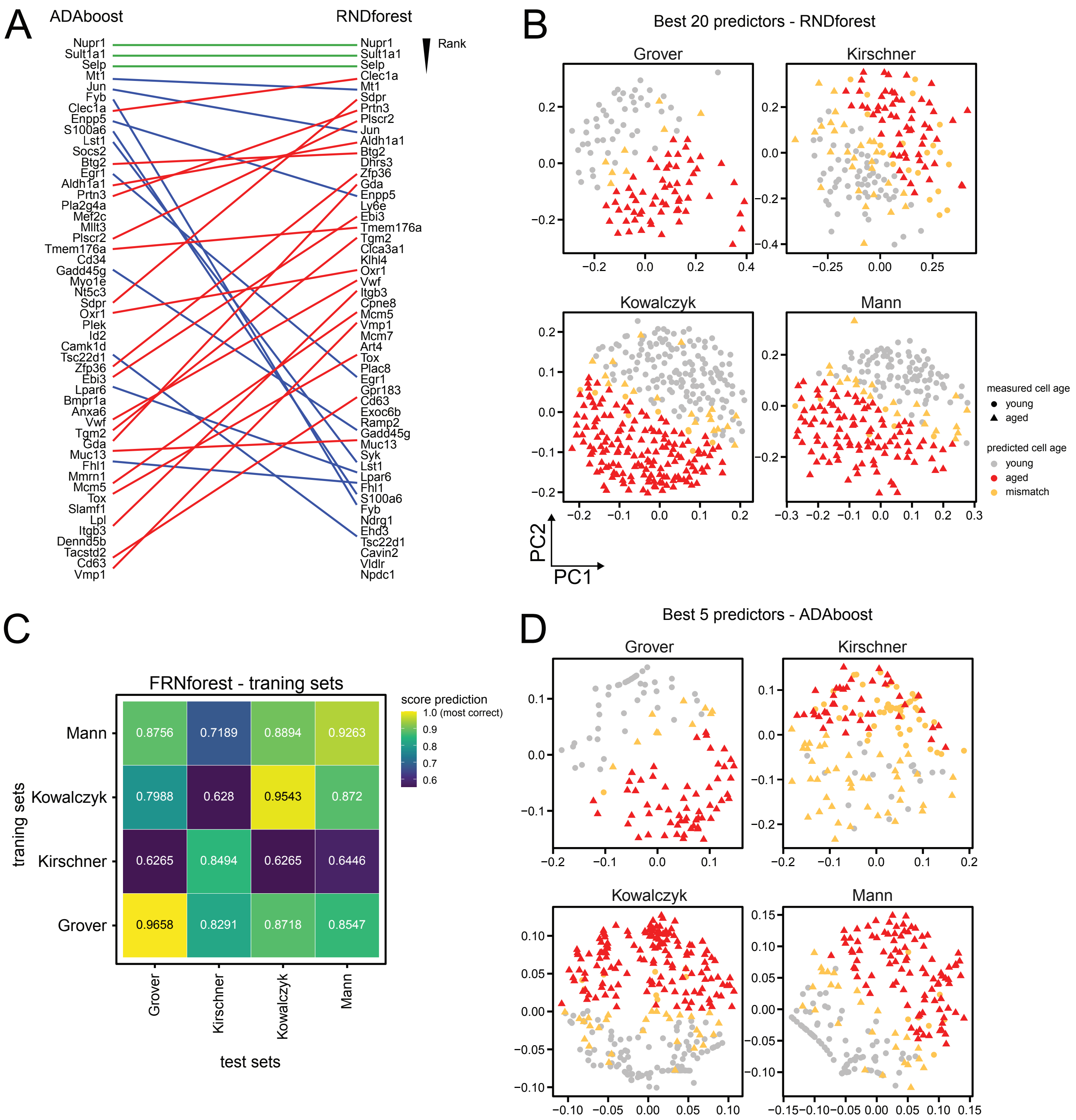

### Supplemental Figure 6

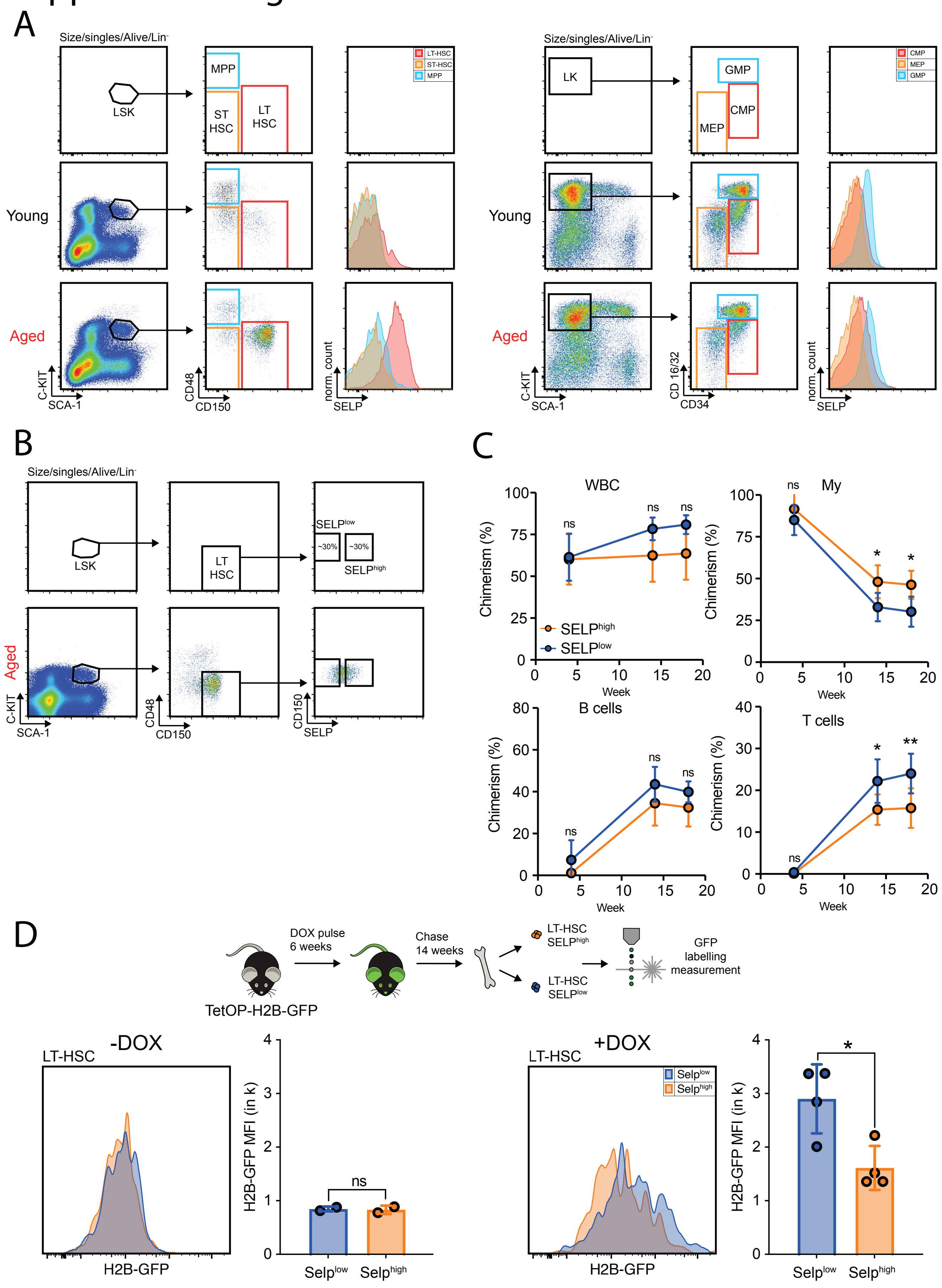
