## Supplemental Text for "A comprehensive transcriptome signature of murine hematopoietic stem cell aging"

**Flow Cytometry**

Bone marrow was isolated from the tibia, femur, pelvis, sternum and spine by crushing, and red blood cells were lysed with erylysis buffer. For LT-HSC isolation, lysed bone marrow (BM) was stained with lineage (Lin) markers antibodies (B220, CD3, GR-1, MAC-1 and TER-119) conjugated with Alexa 700, C.KIT-PE, SCA-1(CD117)-BV421 or Pacific Blue, CD48-Alexa 647, and CD150-PECy7 (all from Biolegend). For Selp HSC isolation, cells was stained with Lin-Alexa 700, C-KIT-FITC (Biolegend), SCA-1-BV421, CD48-Alexa 647, CD150-APCFire 780 (Biolegend), SELP-PE (eBioscience).

For SELP expression on LSKs, BM was stained with Lin-Alexa 700, C-KIT-PE, SCA-1-BV421, CD48-Alexa 647, CD150-BV605 SELP-PE (Biolegend). For SELP expression on HSPCs, lysed bone marrow was stained with Lin-Alexa 700, C.KIT-APC (Biolegend), SCA-1-BV421, CD34-FITC (BD Biosciences), CD16/32-PECy7 (eBioscience) and SELP-PE. For H2B-GFP mice, lysed bone marrow was stained with Lin-Alexa 700, C-KIT-PE, SCA-1-BV421, CD48-Alexa 647, CD150-PECy7, SELP-BV605. For white blood cells chimerism, blood samples were lysed with erylysis buffer and were then stained with CD45.1-PB, CD45.2-PE, CD3-APC, B220-Alexa 700 and MAC-1/GR-1-PECy7 (all from BioLegend).

**Data and Code Availability**

Data generated for this study and collected data is available at Gene Expression Omnibus (GEO). Studies accession numbers used for the analysis are provided in Table S1. Original RNA-sequencing data are deposited under the accession number GSE128050.

We provide two resources. Firstly, we provide a friendly use code-free WEB tool site

(<http://agingsignature.webhosting.rug.nl/>) where users can run their own enrichment analysis and explore the Aging List (AL) and/or Aging Signature (AS). The second resource regards scripts and data analysis, which can be found at GitHub (<https://github.com/LeonidBystrykh/data-for-manuscript>). It contains source files for differentially expressed data (meta and re-analyzed from all used publications). The scripts show major steps in assembling Aging List, Aging Signature. Two applications, find the gene on the list, and check for the enrichment of the set are also provided.

**Data Acquisition**

Pubmed was screened for the following combination of search keywords: “hematopoietic stem” AND (“cell” OR “cells”) AND (“ageing” OR “aging”). The retrieved list was checked for the availability of GEO Datasets. Those papers which also contained original GEO submissions were selected, and further screened for presence of gene expression data in young and physiologically aged mice. In addition, we manually added supplementary information^1^, for which we do not know any publicly available data submission.

**Meta-analysis**

All data for differential expression (DE) for meta-analysis studies were retrieved from provided DE tables from manuscripts and reformatted from files provided by authors and used “as is”.

**Differential expression re-analysis**

Three types of expression data were used: expression micro-arrays, bulk RNA-seq, and single-cell RNA-seq. For all re-analyzed sets, only samples regarding young and aged samples were taken into consideration. No batch effects were assumed or taken for correction.

**Micro-arrays analysis**

Raw data was downloaded from Gene Expression Omnibus (GEO) using available GEO number and analyzed for DE genes using a similiar script structure from online GEO2R (<http://www.ncbi.nlm.nih.gov/geo/geo2r/>). Samples were log2 transformed and quantile normalized using limma R-package. Lists of DE genes were cut by adjusted p-value at 0.05.

**Bulk RNA-seq analysis**

FASTQ files were downloaded from GEO. Following, sequenced reads were aligned to the mouse reference genome (GRCm38, GENCODE M13) using STAR 2.7.0f^2^. Strand specificity was determined using Picard Tools 2.18.0. For differential expression analysis edgeR 3.24.3^3^ was used. Genes with average read count <5 were filtered out; "upperquartile" method was used for library normalization; data dispersion were estimated using function estimateGLMRobustDisp(), significance of differential expression was adjusted by BH method, cut-off values for DE analysis were set by FDR < 0.05.

**Single-cell RNA-seq analysis**

Data for single cell RNA-seq was downloaded as provided by authors from GEO, preprocessed using Scater package ^4^, outliers and cycling cells were removed. The remaining cells were analyzed by Seurat package by comparing two groups of cells, young and aged as provided by authors. No further subgrouping of data was considered. As above, DEG lists were cut by FDR < 0.05.

**Machine learning algorithms and age group prediction**

Machine learning analysis was done in python using custom scripts. RandomForest and ADAboost tools were used from scikit-learn (a.k.a. sklearn) python package as described in sklearn tutorials. Normalizations, scaling, PCA, MDS and other transformations of the single-cell expression data were done with all corresponding tools from the same sklearn package. The script is provided in the dedicated GitHub.

**Coefficient of Variation of consistent subseted datasets**

First, we extracted all gene names from the re-analyzed Aging List (AL) for consistency >1, yielding approximately 1000 genes. For a set of 12 re-analyzed DE lists we generated series of all possible combinations starting from 1 list, and ending with 11 lists (out of 12 in total). Through all permuted combinations we counted mean frequencies of citations for every predefined gene name, and also CV (stdev/mean)

for the same genes. The scatter plots represents the 1000 genes by their mean consistency rank (x-axis) and CV (y-axis).

**Radial plot of individual studies distances to AS**

Degrees of similarity on the Figure 2D (radial plot) were calculated as follows.

we estimated relative frequency (RF) – the number of overlapping genes to a given studie to the AS –The distance is then estimated as:

$$dist=1-RF$$

Node sizes represent the total number of a DE gene list in each data set.

**Resolution modelling**

Resolution – ranging from 0 to 1 - was calculated as the fraction of the consistency genes of subseted studies given a number of studies. Assuming 2 publications, then the gene of interest is either present once, not at all, therefore the resolution of the score is 1/2. The resolution can be written in general terms as:

$$\mathrm{resolution}=1-\frac{1}{n}$$

n = number of studies

**Rank Score**

Rank score is calculated for any particular gene list using simple counting algorithm.

To a given study, genes with consistency score > 1 ( reproducible gene) are summed, otherwise (consistency score < 1, non-reproducible) it is penalized with a score of -1. Thus, the sum of all gene scores is taken across entire list as follows:

$$RS=\sum_{n=1}^{length(test)} \left\{ \begin{aligned} {Sc}_{n} {Sc}_{n}>1 \\ -1 {Sc}_{n}=1 \end{aligned} \right.$$

Test = test gene list

Sc = score: number of times gene was found in all lists (consistency).

**Discrete enrichment tool (dGSE)**

Independently from RS, dGSE provides a score, which is based not just on presence of overlapping genes, but also takes into consideration the order of appearance based on a ranked list. Similarly to GSEA enrichment ^5^, it also utilizes a concept of counting genes from a reference list – in this case AL is taken as such – on top of the test gene list. The test gene list subjected for analysis is first ranked by p-value, most significant on the top. Next it counts the genes from the AL found in discrete subsets of the analyzed set (therefore named discrete gene set enrichment). The subset sizes of the full list are always equal to the sums of sizes of the subgroups in the Aging Signature (beginning from most frequent genes to the least frequent). In our particular case it will be 1 (for *Selp*),1+2 (for *Selp* and [*Mt1*, *Nupr1*]), 1+2+2, 1+2+2+3 and so on.

This analysis comes in two versions. One is calculating enrichment score for the defined window size.

The enrichment score is calculated per fixed window of AS size, 220 genes for re-analysed data. In order to access the overall Enrichment score (EnSc) we firstly determine the number of “observed” and “expected” genes. Observed genes are the number of overlapping genes between the AS and the test set. Expected genes, is defined as the fraction of genes present on the test set and the AS relative to AL (which is 220/5443 (genes in AS divided by AL). Once observed and expected genes are computed, they are subtracted and normalized to the total number of reported genes, AL, and the result is given as %. The following formula shows the expression used:

$$\mathrm{EnSc}=\frac{length\left( AS\cup test \right)- \frac{length\left( test \right)\times length(AS)}{length (AL)}}{length(test)}$$

Test = test gene list

AS = aging signature genes

AL = long list genes

Length(test) = full length of gene list in the test also found in AL

For graphical representation of the EnSc, we perform the same calculation but instead of performed the score calculation for fixed window (AS, 220 genes), we calculate the EnSc for all groups. Thus, the line demonstrate the cumulative EnSc and the red bars at the bottom demonstrate the position of the genes present in the test set in reference to the ranked AL. It gives a visual impression how well any particular set is enriched for AS.

**Quantification and Statistical Analysis**

For figures which statistical analysis was applied, data is presented as biological replicates as mean of *N* events, exact *N* values are provided in each figure legend. For experimental data, data was analyzed using the two-tailed, unpaired *Student’s t test* function or RM two-way ANOVA with Sidak’s multiple comparison test in Prism (GraphPad) for the comparison of two or more groups. Acceptance for *p*-values was set at 0.05 and the significance of each test is provided in each figure legend.

**Supplemental Figure Legends**

**Figure S1 - Related to Figure 1**

A) Mouse age distribution used in all selected studies. Individual dots represent the age (in months) of each age groups from all studies; box plot and violin plots show the distribution of the age groups. Young mice had a median age of 3.03 ± 1.04 (SD) months and aged mice a median age of 22 ± 3.43 months.
B) Re-analyzed studies cluster by sequencing platform. PCA of normalized and centralized re-analyzed datasets. PC1 and PC2, which are responsible for the majority of the variation, are shown. Studies are color labelled according to the sequencing platform used.
C) Correlation between fold-changes (FC) found in studies which are present in both meta and re-analyzed analysis. Dots represent individual DEG genes found in both analysis, and lines represent linear curve fitting and its R^2^ is shown.
D) Combined number of consistently reported DEGs across studies. Dots represent the number of Differentially Expressed (DE) genes (y-axis) and in how many studies they were reported to be differentially expressed (x-axis). Lines represent the curve fitting to each analysis. Purple and green dots/line indicate meta-analysis and re-analysis, respectively.

E/F) Heatmaps showing the pair-wise comparison of studies in meta and re-analyzed analysis. Numbers indicate the number of overlapping genes each study has with another study. The color of each square represents the number of overlapping studies. Fraction of total overlaps represents the percentage of reported genes which overlapped with any other study.

G) Bar plot depicting the number of all reported DEGs. Purple and green bars demonstrate the number of genes which are reported in ≥ 4 studies in meta and re-analysis, respectively. Gray bars indicate the number of DEGs which were found in ≤ 3 studies.

**Figure S2 – Related to Figure 2**

A) Gene Ontology enrichment analysis of all genes in the aging signature. All genes, regardless of their directionality were used for GO analysis. Represented are the top 10 terms from unfiltered GO “molecular function” category. Numbers in each bar demonstrate the number of genes found per term.
B) Pathway analysis of AS genes divided by directionality. KEGG and REACTOME significantly enriched terms are displayed. Numbers in each bar demonstrate the number of genes found per term. Red and blue bars represent GO terms found to be significantly enriched in upregulated and downregulated AS genes, respectively. Terms were selected based on a FDR < 0.05 cutoff.
C) Schematic representation of how associated GO terms were selected. Upper panel: All associated GO terms for individual AS genes were collected from UNIPROT, termed AS GO terms database (upper panel). Following we overlapped genes which contained similar GO terms and plotted the number of genes (x-axis) which shared similar GO terms (y-axis). The large majority of GO terms have just a small number of AS genes.
D) From the AS terms database, we performed keyword search for different HSC aging mechanisms such as “membrane”, “inflammation”, “histone” and “cell cycle” (x-axis). The number of GO terms found to be associated with each keyword is shown (y-axis).

**Figure S3 – Related to Figure 3**

A) Isolation of LT-HSCs. Upper panel shows a schematic representation of the FACS gating strategy for isolation of LT-HSCs. Middle and lower panel show the population frequencies for young (middle panel, grey) and aged (lower panel, red) mice.
B) RNA content per LT-HSC cell from young and aged mice (n = 8 per group).
C) Quantification of MFI of SYTO of sorted LT-HSC cells derived from and young (grey, n = 120 cells) and aged (red, n= 139 cells).
D) Overlap of total number of chromatin accessible sites found in young and aged HSCs. Each group is comprised of peaks which were called in both biological replicates per group (grey, young and red aged).
E) Broad peaks of differentially accessible sites (peaks) between young and old HSCs measured by ATAC-seq. Dot colors demonstrate peaks either in genomic regions overlapping with coding (red) or non-coding (blue) annotations. Peaks with negative FC values are significantly more accessible in young HSCs whereas peaks with positive FCs values are more significantly accessible in aged HSCs. Gene symbols annotate peaks which overlap with gene bodies from some AS genes.

For panels D, E, ± SD is shown. **** p < 0.0001.

**Figure S4– Related to Figure 4**

A) Enrichment scores for different transcriptomic sets. dGSE analysis and RS for the upregulated genes in the de Magalhães signature (blue) and different TGF-β pathway genes (green). Left demonstrates the enrichment scores for different de Magalhães and TGF-β data sets together Wong data set (purple, as a negative control) and the HSC aging signature itself (red, positive control). Right demonstrate the scores of these data sets. Numbers indicate the data set with legend below the scores.
B) Separation of young (circles) and aged (triangles)single-cell transcriptomics and their respective AS score. From Kirschner (80 young and 85 aged cells), Kowalczyk (152 young and 174 aged cells) and Mann (95 young and 120 aged cells, individual cells were separated by PC1 (x-axis) and PC2 (y-axis) and color-coded according to their individual aging signature score. Left column display only young cells; middle column shows just aged cells and right column shows both populations. Cells are colored score from low (blue) to high (yellow) AS scores. Violin and boxplot demonstrate the AS score (y-axis) distribution of the young (grey) and aged (red) cells per study.

**Figure S5– Related to Figure 5**

A) Comparison between rank genes using 2 different machine learning algorithms. Gene symbols from genes rank by their best capability of separating young and aged HSCs. Left column shows the best ranking genes using ADAboost and the right column genes for RNDforest. Green lines demonstrate the genes which score similarly in both algorithms and blue and red lines demonstrates de different scores that genes have depending which algorithm was used.
B) Young (circles) and aged (triangles) single-cell were separated by PC1 (x-axis) and PC2 (y-axis) and color-coded according to the match between the measured aged of the cells and the predicted aged measured by the Random Forest algorithm(RNDforest) (grey for young cells; red for aged cells and orange for mismatched cells).Right)Machine learning scores varies depending on which training set is used. Heatmap from RNDforest training depicting different overall scores for different scRNA-seq sets used. The overall score is color-coded from blue (lower scores) to yellow (higher scores). Sets on the x-axis (training sets) were used to train the algorithm and the following sets on the y-axis (test sets) were scored according to training.
C) Machine learning scores varies depending on which training set is used. Heatmap from RNDforest training depicting different overall scores for different scRNA-seq sets used. The overall score is color-coded from blue (lower scores) to yellow (higher scores). Sets on the x-axis (training sets) were used to train the algorithm and the following sets on the y-axis (test sets) were scored according to training.
D) Using the best 5 gene predictors for population separation in different scRNA-seq sets. Young (circles) and aged (triangles) single-cell were separated by PC1 (x-axis) and PC2 (y-axis) and color-coded according to the match between the measured aged of the cells and the predicted aged measured by the Random Forest algorithm(ADAboost) (grey for young cells; red for aged cells and orange for mismatched cells).

**Figure S6– Related to Figure 6**

A) Isolation of different hematopoietic populations. Left upper panel shows a schematic representation of the FACS gating strategy for isolation of LT-HSC, ST-HSC and MPP; Right upper panel depicts the gating for GMP, CLP and MEP. Middle and lower panels show the population frequencies for young (middle panel, black) and aged (lower panel, red) mice.
B) Isolation of SELP^low^ and SELP^high^ LT-HSC. Similar FACS gating strategy was used for LT-HSC **(see above)**. SELP^low^ and SELP^high^ fractions of LT-HSC population were defined as the lower 30% and the higher 30% SELP-expressing cells.
C) Lineage output at 18 weeks post-transplant aged Selp^low^ (blue)and Selp^high^ (orange) LT-HSCs produced more myeloid cells (n = 6-7 animals/group). Each dot represents a mouse in each group(White blood cells) indicate total donor chimerism while My (myeloid) cells, B and T cells cells are shown as a percentage of this compartment.
D) SELP expression is associated with increased HSC cycling. The left panel shows the experimental approach. Right panels show H2B-GFP expression in DOX treated mice (n = 4 mice for each group) and control mice that were not treated (n = 2 for each group). Right upper panel shows the H2B-GFP expression in LT-HSCs subdivided in SELP^low^ (blue) and SELP^high^ (orange). The far right panel shows that Selp^low^ LT-HSCs preferentially retained the GFP label, which is diluted in Selp^high^ LT-HSCs.

For panels D, E, ± SD is shown. * p < 0.05.

**Supplemental Table Titles**

**Table S1: Overall information of collected transcriptome studies used in this study**

**Table S2: Gene symbols for meta and re-analyzed aging signature and aging list**

**Table S3: Gene Ontology (GO) and pathway analysis of re-analyzed aging signature**
